# Group II Introns in Archaeal Genomes and the Evolutionary Origin of Eukaryotic Spliceosomal Introns

**DOI:** 10.1101/2024.12.10.627823

**Authors:** J. S. A. Mattick, S.-B. Malik, C. F. Delwiche

## Abstract

A key attribute of eukaryotic genomes is the presence of abundant spliceosomal introns that break up many protein-coding genes into multiple exons and must be spliced out during the process of gene expression. These introns are believed to be evolutionarily derived from group II introns, which are known to be widespread in bacteria. One prominent hypothesis is that the spliceosomal intron arose after the endosymbiotic origin of the mitochondrion, as a consequence of transfer of genes containing group II introns from the organelle to nuclear genome; in this model, transfer of group II introns into the ancestral eukaryotic genome set the stage for evolution of the spliceosomal form. However, the recent discovery and sequencing of asgard archaea — the closest archaeal relatives of extant eukaryotes — has shed significant light on the composition of the early eukaryotic genome and calls that model into question. Using sequence analysis and structural modeling, we show here the presence of group II intron maturases in the genomes of Heimdallarchaeia and other asgard archaea, and demonstrate by phylogenetic inference that these are closely related to both eukaryotic mitochondrial group II intron maturases and the spliceosome protein PRP8. This suggests that the first intron-containing eukaryotic common ancestor (FIECA) inherited selfish group II introns from its ancestral archaeal genome – the progenitor of the nuclear genome – rather than from the mitochondrial endosymbiont. These observations suggest that the spread and diversification of introns may have occurred independently of the acquisition of the mitochondrion. To better understand the context for intron evolution, we investigate the broader occurrence of group II introns in archaea, identify archaeal clades enriched in group II introns, and perform structural modeling to examine the relationship between the archaeal group II intron maturase and the eukaryotic spliceosome. We propose a model of intron acquisition and expansion during early eukaryotic evolution that places the spread of introns prior to the acquisition of mitochondria, possibly facilitated by the separation of transcription and translation afforded by the nucleus.

## Introduction

Introns are intragenic sequences that must be removed from the gene sequence with a dedicated splicing machinery before the protein encoding portion of the RNA may be translated^1^. They are pervasive in most eukaryotic genomes in the form of spliceosomal introns, but the origins of their entry and spread in early eukaryotes remains opaque^2^. Intron splicing is critical for viable gene expression in almost all eukaryotic species, and this near-universal distribution indicates that the spliceosome traces its evolutionary history to the last eukaryotic common ancestor (LECA)^3^. Introns also allow alternative splicing such that a single gene can yield variant transcripts, and hence have been postulated to contribute significantly to the diversification and complexity of eukaryotes^4–6^, although splicing mechanisms in many protists remain poorly understood^7^. As one of the major information processing innovations that occurred prior to LECA, introns represent one avenue of diversification that emerged in stem eukaryotes, during the transition from the prokaryotic *first* eukaryotic common ancestor (FECA) to the complex organellar cell that LECA was likely to have been (*i.e.*, during the early evolution of the eukaryotic lineage)^8^. Recent discovery and genome sequencing of the asgard archaea are able to provide new insights into the development of intron systems in these organisms, and how they might be adapted into eukaryotic lineages that share a common ancestor with asgard archaea (see section “Archaea in the tree of life”, below, for a discussion of the nomenclature we use here).

Spliceosomal introns are the primary form of intron present in eukaryotes, but they are not the only form^9^. Spliceosomal introns are characterized by the spliceosome, a megadalton-sized core of proteins coupled to small RNAs that act as ribozymes and catalyze the two transesterase reactions required to remove intronic sequences^10^. Many of the spliceosomal proteins are regulatory in nature, but the core protein PRP8 interacts directly with the U5 small RNA to recognize the 5’ splice site and bring it into proximity with the 3’ splice site for trans-esterification^11–13^. There are two major classes of spliceosomal introns: the U2 and U12 intronic systems. These systems both share a catalytic core, and almost certainly share an evolutionary origin in LECA, but differ in kinetics and in gene preference^14^. The U2 system is responsible for 99.5% of the splicing in most eukaryotic systems, but disruption of the U12 system has significant phenotypic consequences in vertebrates^15^.

### Mechanisms of splicing

By contrast, other classes of introns are largely self-sufficient and can be divided into three categories: group I introns are self-splicing RNA sequences that do not require protein chaperones to complete the transesterification necessary for splicing^16^. The predominant mechanism for their spread is thought to be by association with an invasive homing endonuclease sequence, but evolutionarily these proteins are not conserved with the splicing apparatus and can be gained and lost independently from viral or retrotransposon elements^17^. Group II introns are found in bacteria, archaea and bacteria-derived organelles^18,19^. Unlike group I introns, group II introns always carry the machinery necessary for endonuclease-based invasion. These genetic elements are comprised of two components: the first is a coding sequence within the spliced RNA (the “intron”), which codes for a protein called a maturase. The maturase contains domains that assist in forming the splicing “lariat” (a loop of RNA reminiscent of a cowhand’s lariat), have homing endonuclease activity which can create double stranded breaks in uninfected alleles^20^, and confer reverse transcriptase activity which will then copy the entire intron into the break during homologous repair^21^. The second key component of the group II intron structure is a specific secondary structure of RNA that flanks the coding sequence of the maturase, and that catalyzes the actual splicing as a ribozyme^22^. It has been noted previously that there are significant similarities between the structures of the group II ribozyme and the spliceosome intron complex found in eukaryotes^23^: Both utilize a protein (either a maturase or the eukaryotic protein PRP8) to stabilize the ribozyme complex^24^, both form similar RNA secondary structures, including the splicing lariat, and both utilize Mg^2+^ ions as a core component of catalysis (with both being rescued by the same ions when subjected to oxygen-sulfur substitution)^25^. Furthermore, structural analysis of the core spliceosome protein PRP8 and the maturase of group II introns provides strong evidence of homology between the eukaryotic spliceosome and these genetic elements, including what appears to be a non-functional reverse transcriptase domain within PRP8 itself^26,27^. Rounding out the catalog of introns, group III introns are a class of self-splicing introns that are only found in euglenoid plastids^28^. These introns form the same splicing lariat as other groups, but do not contain the same structure as group II introns, although they often contain maturases that are closely related to the group II intron maturase. Finally, there are unrelated introns, including those in tRNAs, whose removal is entirely protein catalyzed.

One of the key differences between spliceosomal and selfish introns (groups I-III) is the presence of a mechanism for self-propagation^29^. For group II introns, this is accomplished *via* a reverse transcriptase domain that allows these sequences to function much like retrotransposons, with the caveat that they do not carry their own promoter, and thus must jump into active genes to maintain activity^30^. In addition, unlike retrotransposons, group II introns can remove themselves from the primary transcript they are embedded in *via* splicing, allowing them to function with minimal disruption to the host organism^31^. Furthermore, because their splicing is dependent on both a protein maturase and a strong RNA secondary structure capable of lariat formation, it is very difficult to disrupt or remove the invasive element while retaining the intact coding gene^32^. In principle, this should allow group II introns to spread as pervasively in prokaryotes as spliceosomal introns have in eukaryotes, but this has not been observed^33^. Selective pressures stemming from large populations and small genome sizes are likely capable of removing these sequences from populations^34^, but it is possible that other differences in eukaryotic biology account for the relative success of intron fixation between domains. We surveyed the distribution of group II introns in archaea and bacteria with the objective of understanding the success of these retroelements in both prokaryotic domains and elucidating their role in the evolution of the eukaryotic lineage.

### Archaea in the tree of life

Until recently, the three basal branches on the tree of life were thought to be Bacteria, Archaea, and Eukarya, but identification of Asgard archaea has shifted the predominant view of the phylogenetic tree of life^35–37^. There is strong support for a close relationship between Asgard archaea and eukaryotes, with recent analyses placing Hodarchaeales as the sister group of Eukarya within the larger lineage of Heimdallarchaeia^38,1^. Although they do not refer to monophyletic groups, we will use the informal terms prokaryote, archaea, asgard, and heimdallarchaeia (in lower case) for convenience and clarity. The presence of many genes previously thought to be eukaryote-specific in the asgard archaea has shed some light in the order and mechanisms involved in the origin of eukaryotes, although there remains significant debate about both the order of events and the role that bacterial symbiosis played in it^41^.

However, due to evidence for the current or past presence of mitochondria in all known eukaryotes, it seems likely that the bacterial symbiosis leading to the origin of the mitochondria occurred at some point along the long branch between archaea and eukaryotes^42^, *i.e.*, in stem eukaryotes between the FECA and LECA. Introns have also been traced back to this long branch, with almost all currently known eukaryotes having some form of splicing machinery in their genomes^43,44^. The presence of group II introns in mitochondria has led to the hypothesis that introns entered into the eukaryotic genome through endosymbiotic gene transfer from the early mitochondrion, rather than being present at the FECA^19^.

To test this hypothesis, we developed a reference phylogeny of 289 taxa from archaea, bacteria and eukaryotes (a “tree of life”) using 20 universal ribosomal proteins from the large and small subunits and 19 other universal proteins that were detected in all three domains of life across multiple taxa with a BLAST threshold of 1E-5 and were reciprocal best hits to query proteins.

535 taxa were chosen for the database to ensure broad taxonomic representation and minimize bias toward any one lineage by overrepresentation. Taxa that had all 39 of these proteins were included, with the reasoning that any proteomes that were lacking these universal proteins were likely to be incomplete. The selected proteins were then aligned and concatenated, and a maximum likelihood tree was inferred to describe their relationship (**Figure 1**). Group II intron maturase protein sequences were queried against the 535 taxon database, and their presence in genomes was overlaid onto this tree, identifying group II intron sequences spanning the tree of life from bacteria to multicellular eukaryotes (**Figure 1**). Our results are consistent with previous studies that show group II introns as primarily mitochondrial in eukaryotes, and mainly found in methanogens in the archaea^27,44^. However, our work identifies group II introns distributed across asgard archaea, including heimdallarchaeia, the closest relatives of eukaryotes. This suggests that the common ancestor of eukaryotes and Asgard archaea likely also encoded group II introns in its (host) genome prior to the acquisition of the primary mitochondrial endosymbiont.

**Figure 1.**
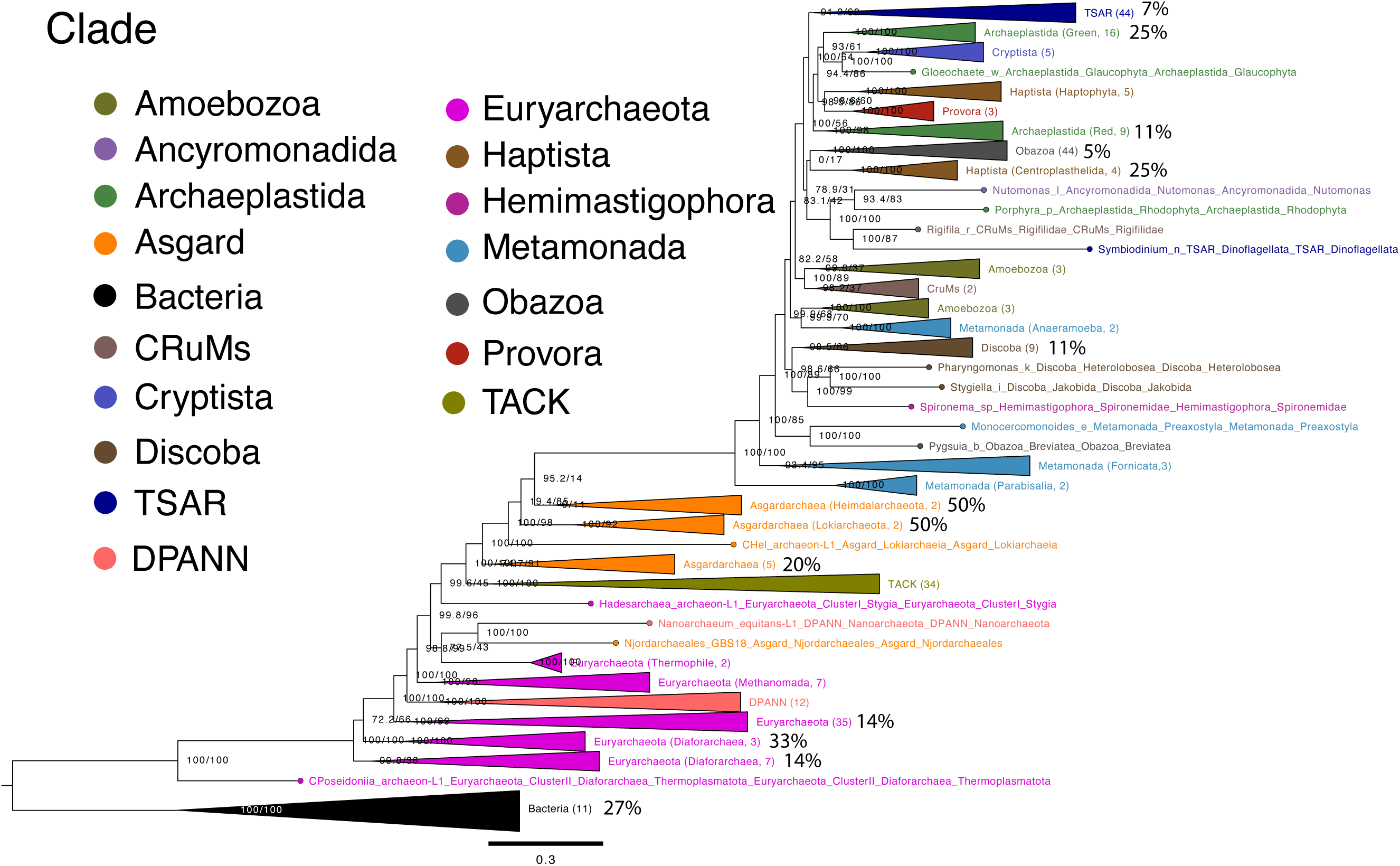
Group II intron distribution across the tree of life. A reference phylogeny based on a concatenated dataset of 39 ribosomal protein and other genes from 289 taxa; tree found by maximum likelihood. Group II introns were identified from all three domains of life using representative taxa (**Supplemental Figure 1**). The percent of taxa containing group II introns identified in each clade are displayed, while the total members of each clade are shown in brackets. Supergroups that are only monophyletic at the order or phylum level are also specified in brackets. Group II introns persistent in eukaryotes are exclusively located in the mitochondria and are primarily found within Archaeplastida and Rhizaria. Archaeal Group II introns are common in methanogens and Asgard archaea. Bacterial group II introns were identified in *Chlorobium* and *Blastopirella*, although because they are serving here only as an outgroup, Bacteria are poorly represented in this tree and this does not capture the breadth of their distribution in that domain.

Our phylogenetic analysis includes diverse eukaryotic, archaeal and bacterial taxa, and was used to determine the distribution of introns across the tree of life. The analysis is based on a relatively short alignment of amino acid sites and highly diverged taxa, and consequently is not expected to capture all of the features observed in more densely sampled and taxonomically focused phylogenies^38^. Our analysis places the eukaryotic root within the asgard archaea, with Hodarchaeales as the sister taxon to eukaryotes (among currently known organisms). It also recapitulates major archaeal supergroups TACK and DPANN as monophyletic. Surprisingly, euryarchaeotes are not monophyletic, but their placement with DPANN as the earliest diverging archaeal groups is consistent with other analyses. The lack of monophyly could well be an analytical artifact. The eukaryotic portion of the phylogeny shows many well recognized monophyletic groups, but has low bootstrap support for most deep-branching features, and some taxa are anomalously placed. This likely represents a complex mixture of phylogenetic artifacts (including from compositional bias and rate heterogeneity) and the complex genomic history caused by endosymbiosis. The tree agrees with other recent analyses placing eukarya within asgard archaea, and permits us to examine the distribution of Group II introns across the phylogeny.

### Group II introns across the tree of life

Group II introns were first discovered in plant mitochondria and then in free-living bacteria, although they were soon also found to be present in archaea^19^. Due to their absence from eukaryotic nuclear genomes and their seemingly limited distribution in archaea, they have often been interpreted as having originated in bacteria with independent lateral transfer into both eukaryotes and archaea^19,27^. However, Vosseberg et al. found that many asgard archaeal genomes do contain group II introns^46^, which raises the possibility that FECA encoded group II introns in its genome prior to the divergence of asgard archaea and eukaryotes (alternatively, they may have undergone multiple transfers between bacteria and archaea, but see below). As of yet, analysis of of these intron maturases have been unable to resolve whether or not these Asgard group II introns are were present in LECA, which raises the question of just how pervasive group II introns are in the archaea, as well as the details of their biology. As group II introns require both a functional maturase and a robust RNA secondary structure to remain active, we sought to characterize group II introns across archaea at both the protein and RNA level.

Using the database and relationships established in **Figure 1**, group II intron maturases were identified *via* PSIBLAST seeded with several known eukaryotic group II introns. Initial results were filtered at a permissive e-value of 1e-1 to identify all reverse transcriptase proteins in the database. This was further filtered *via* the identification of specific maturase domains (Intron_maturas2, GIIM) using PFAM with a confidence threshold of 1e-20. The genomic and metagenomic contigs from which the protein sequences were translated were examined with RFAM for the identification of group II intron RNA structures, and protein sequences were filtered to only include those whose RNA also contained high confidence (1E-10) group II intron structures. This subset of group II intron protein sequences was aligned, trimmed (**Supplementary Figure 1**), and used to infer a maximum likelihood phylogenetic tree (**Figure 2**). The resulting tree was rooted arbitrarily **(Figure 2, Supplementary Figure 2)**, and shows that most mitochondrial sequences form a monophyletic group. Our analyses unambiguously document the presence of functional group II intron maturases in the genomes of multiple Asgard archaea, including Heimdallarchaeales, Lokiarchaeia, Thorarchaeia and Njordarchaeales. Group II introns are found in all the archaeal supergroups, although only within rather restricted taxonomic groups in each of them. Although horizontal gene transfer may well have played a role in their evolution, it appears to have been a common and ongoing process throughout archaea comparable to other forms of genetic exchange, not an exceptional event that occurred at the origin of eukaryotes as has often been inferred. At the very least, if the distribution is to be explained by gene transfer, there have been many such transfers, and transfer directly into the FECA genome would not be surprising. Our analysis of both protein domains and RNA secondary structures (**Supplementary Figures 3,4**) suggests that Asgard archaeal group II introns remain catalytically active with intact secondary structures capable of transesterification catalysis. Given the close relationship of these taxa to eukaryotes, this suggests that group II introns were present and functional in FECA prior to the acquisition of the mitochondrion, which suggests that their expansion into the spliceosomal system during early eukaryotic evolution was not a matter of acquisition but rather of diversification. Furthermore, the bacterial taxa in which we identified group II introns (*Bdellovibrio* and *Chlorobium*) are a predatory and a sulfur oxidizing bacterium, respectively; neither of these has been discussed as a contributor to FECA even in hybrid-origin scenarios.

**Figure 2.**
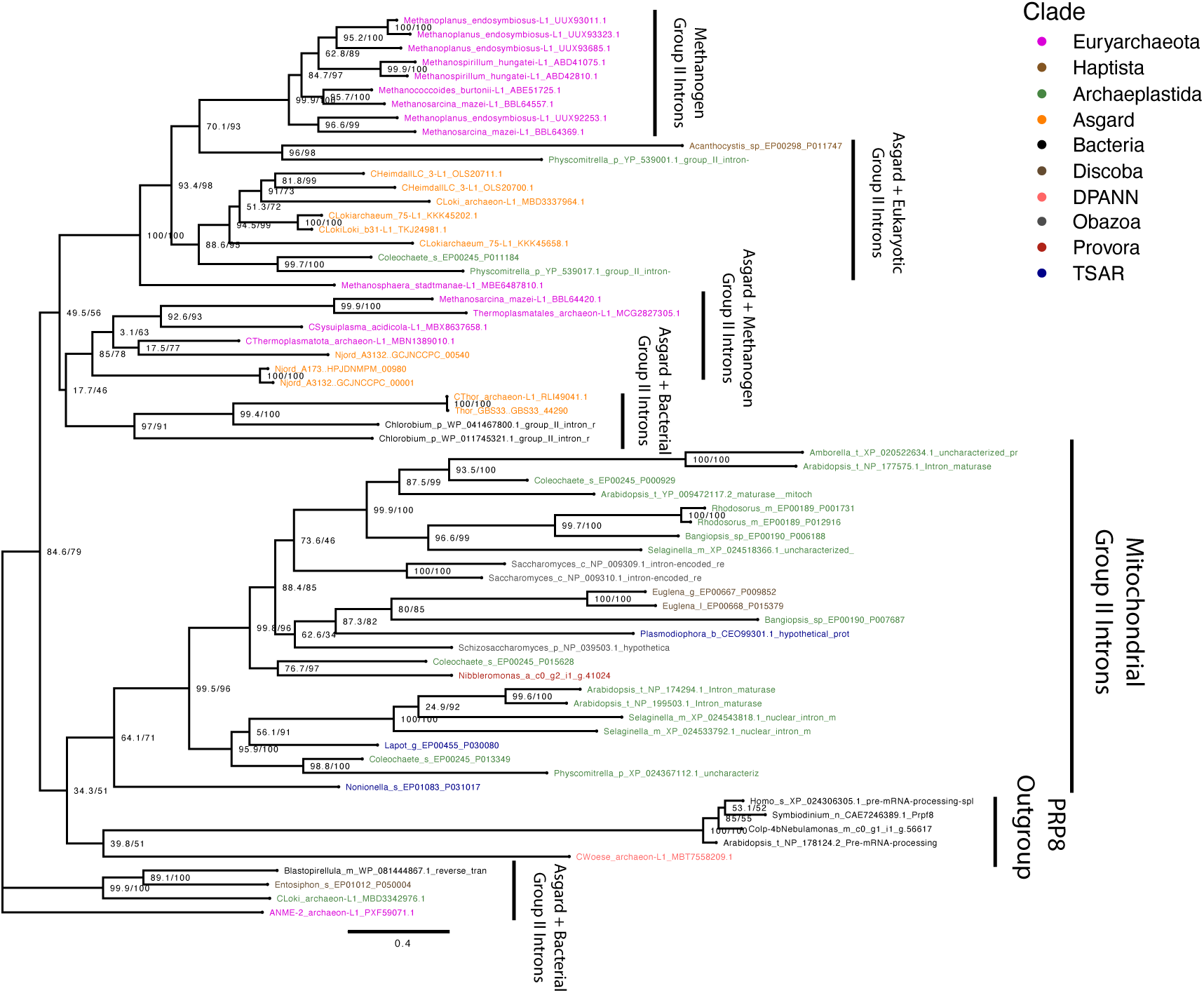
Group II introns similar to eukaryotes were present in Asgard archaea independent of bacterial symbiosis. Representative group II introns were queried using PSIBLAST against representative eukaryotic, archaeal, bacterial and viral databases. Threshold for inclusion was set at 1e-1 and the search progressed for 10 iterations. Results were then filtered for proteins that contain the characteristic group II intron maturase domain. The resulting tree was arbitrarily rooted and indicates that bacterial group II introns appear to be more similar to group II introns already present in the Asgardarchaea than to eukaryotic organellar group II introns. Group II introns were also detected in DPANN, euryarchaeotes and TACK.

We were not able to determine from Alphafold structures of asgard and bacterial group II intron maturases which of the two structures is more similar to PRP8. We attribute this to the fact that group II intron maturases are substantially constrained by their function, resulting in similar structures across all taxa. One leading scenario for the acquisition of the mitochondrion focuses on symbiotic metabolic partnerships involving hydrogen reduction. Our analyses did not identify any such organism with group II introns, although given the patchy distribution of intron sequences across the tree, future identification of an example is always possible. Neither, however, is there evidence that the mitochondrial group II intron sequences are of proteobacterial ancestry as would be predicted by hypotheses that infer a mitochondrial origin of spliceosomal introns. Mitochondrial group II intron sequences formed a distinct clade in our analyses and did not specifically cluster with any archaeal or bacterial sequences (**Figure 2**). A few eukaryotic sequences fall outside of the mitochondrial clade and cluster instead with archaea, suggesting that they may be much more recent integrations of group II intron sequences than those in the mitochondrial clade.

### Archaeal distribution and abundance of group II introns

Another important consideration is intron abundance, because (as with any mobile element) copy number can increase *via* intragenomic gene invasions. To determine the abundance of group II introns across archaea, PSI-BLAST was used to query all archaeal species in Genbank with the same filters used to prepare **Figure 2**. The resulting tree, **Supplementary Figure 2**, indicates that group II introns are present in all supergroups of archaea, including many TACK and DPANN archaea. In phylogenetic analysis the asgardarchaeal group II introns are closely related to those of the methanoarchaea, a euryarchaeal group that is quite distant from both eukaryotes and bacteria (**Figure 1**), consistent with the distribution of group II introns in bacterial taxa. We then assessed evidence that these group II introns from archaea are active (*i.e.,* would be predicted to have catalytic activity) by examining contigs containing group II introns using RFAM for canonical group II intron structures. The number of group II introns that appear to contain both an active maturase and an active catalytic RNA secondary structure within each genome was then calculated (**Supplementary Figure 3**). Strikingly, archaea of genus *Methanosarcina* encode many group II introns, while other archaea, including the asgard, generally encode one or two copies in their genomes. When analyzed for abundance per species, introns remain rare in archaea. This is in contrast with the pervasive nature of spliceosomal introns in eukaryotes.

This overall pattern is consistent with the abundance of retrotransposons and other selfish elements in prokaryotes, which are thought to be under significant negative selection and are constantly in an equilibrium of purge and gain from their host genomes^47^. This suggests that group II introns have a negative phenotypic effect on the host organism (as do retrotransposons), despite their ability to self-splice from a transcript to leave an intact coding sequence^48^. One possibility is that the coupling of transcription and translation in prokaryotes means that there is not enough time for splicing to consistently occur before ribosomes attempt read through the gene that has been inserted **(Figure 3a)**. Further evidence would be required to confirm this hypothesis, but it would be consistent with the eukaryotic phenotype, which is able to tolerate massive intronic intrusions *via* the decoupling of transcription and translation in the nuclear membrane. It is also compatible with the hypothesis that the large population sizes of bacteria and archaea (compared to most eukaryotes) favors small genomes and genomic efficiency^34^.

**Figure 3.**
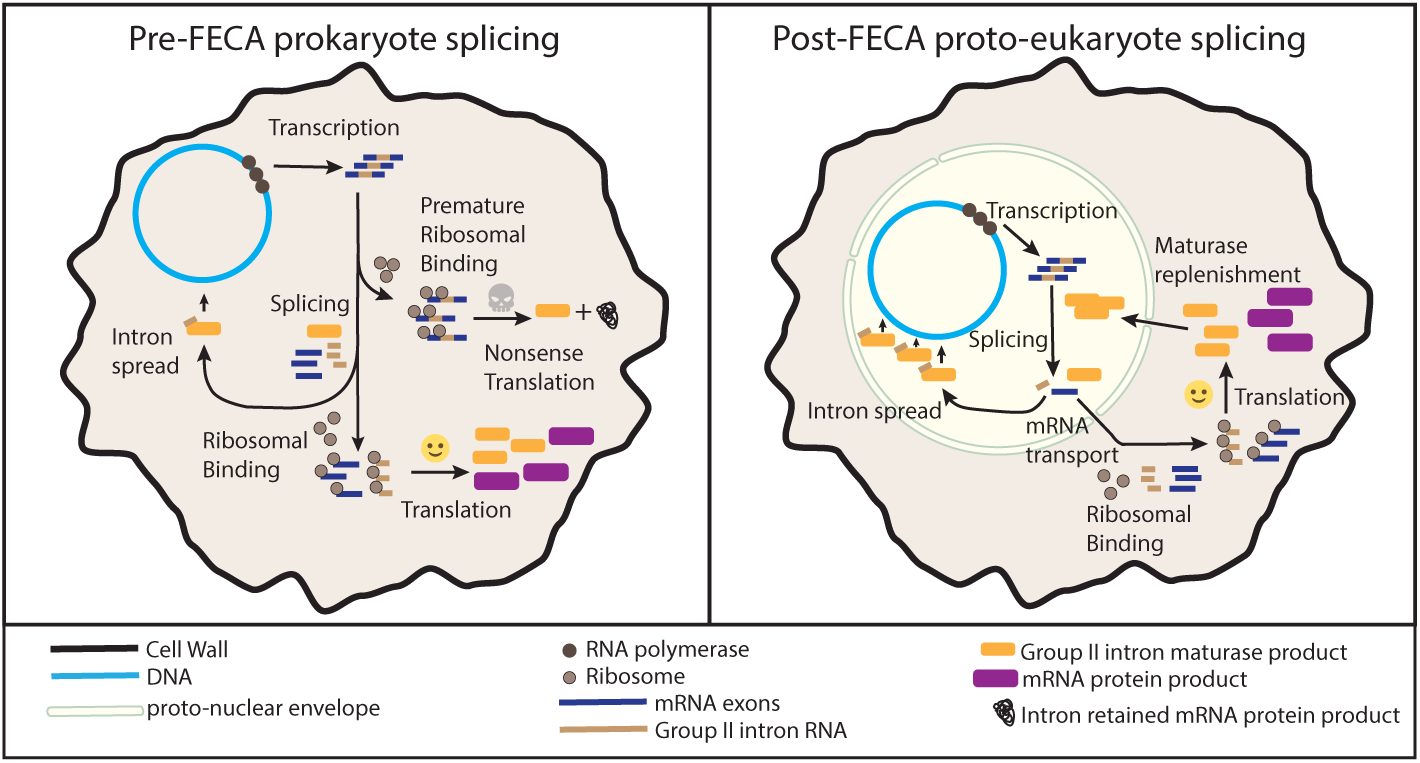
Model for intron spread in the early eukaryote. In this model of intron spread during early eukaryogenesis, the group II intron RNA sequences that are the common ancestors of the U2 and U12 spliceosome, and the maturase that is encoded are spliced into transcripts. In the pre-FECA model, there is no separation of transcription and translation such that ribosomes are competing with the maturase population for splicing. If the ribosomes attach to translate before the maturase can splice the transcript, the products will be intron-inclusive and likely nonfunctional. Thus, every group II intron insertion back into the genome comes at significant cost to the organism in the form of partially incomplete proteins. In contrast, the proto-eukaryote with some form of nuclear envelope has separation between transcription and translation, allowing for maturase access to the intron prior to ribosomal exposure. This allows for protein synthesis to complete only on spliced and exported transcripts, removing much of the cost of intron inclusion. Since the maturase is required to be within the nuclear envelope for this functionality, it is also able to rapidly spread introns throughout the nascent genome without significant fitness costs to the organism.

### A model for intron spread in the nascent eukaryote

The widespread presence of group II introns in archaea suggests that the early eukaryotic nuclear genome was almost certainly exposed to these genetic elements, but the patchy distribution and low abundance of these elements indicates that they are rarely abundant, and were probably not so in the early stages of divergence from other asgard archaea. Interestingly, we identify no group II introns in eukaryotic nuclear genomes. Previous work has shown that group II introns expressed in the nucleus are capable of successful self-splicing in the cytoplasm^49^, although the contained transcript is poorly translated, and often subject to nonsense mediated decay. This can be attributed to the fact that successful splicing of the group II system requires that the transcript itself form a specific RNA lariat for catalysis. This has been separated in eukaryotic systems from translation. In the cytoplasm of archaea and bacteria that contain group II introns but do not have a nucleus, the binding of the maturase protein and the completion of the splicing reaction must compete with ribosomal complex binding and translation. In **Figure 3**, we outline a model for intron spread during early eukaryotic evolution, driven by the cost of retained introns in an unseparated transcriptional and translational system, and independent of the mitochondrion. Pre-FECA, the group II intron system would have produced transcripts that must be spliced while being actively exposed to the ribosome.

Ribosomal binding and translation prior to a successful splicing event not only produce intron-retained proteins, but also likely prevent future splicing as more ribosomes bind for further translation. This imposes an inefficiency on the invaded transcript, and creates the risk that intron-retained proteins may misfold or aggregate. However, in early eukaryotic evolution the innovation of the nucleus to separate transcription and translation would have been able to rescue this inefficiency by removing the ribosome from the splicing compartment. The reduced burden of group II introns on the nascent eukaryote would then have allowed them to be retained, and could have spread throughout the genome with little cost to fitness. Another major innovation of the eukaryotic spliceosome, in contrast to a group II intron system, is the ability to recruit small RNAs that re-capitulate the group II secondary structure for ribozyme formation in splicing. This allows trans-splicing to remove intronic sequences without conservation of the secondary structures in the transcript itself, and decouples spliceosome expression from the transcription of any given gene. With the reduced burden of splicing efficiency in the early eukaryote, the ancestral group II intron sequence could spread rapidly within the nascent nuclear genome. Once there was a significant intron burden within the genome, natural selection would act to rescue inefficient splicing. The implications of this are that the evolution of the spliceosome was driven by the need to trans-splice these sequences to rescue the nascent eukaryote from the mutational burden and potential disruption of large numbers of group II secondary structures. This helps explain why mitochondrial group II introns can be obtained through gene transfer but are rapidly lost when they enter the eukaryotic nucleus. It is also consistent with the fact that although many mitochondrial genes have been transferred to the nucleus, and many other selfish elements can persist in eukaryotic genomes, group II introns have never been detected there.

## Conclusions

We show that group II introns are widespread in the asgard archaea, and hence were likely present in FECA, or at least a close relative of it. The widespread occurrence of group II sequences and concordance between the group II intron phylogeny and the organismal phylogeny (in the sense that monophyly of sub-groups is maintained) suggests that vertical inheritance is important, and likely the predominant mechanism for intron inheritance within this clade, although we cannot exclude the occurrence of some horizontal gene transfer as well. Furthermore, we show that group II introns are not enriched in abundance in asgard archaea relative to other archaeal species, which indicates that intronic expansion and the origin of the spliceosome occurred after FECA had diverged from other asgard archaea, but need not have been correlated with the origin of the mitochondrion. In this context, we propose a model whereby intronic expansion required the separation of transcription from translation *via* the nucleus to overcome the negative selection on these species independent of bacterial lineages, and that the mitochondrion was acquired after the evolution of the spliceosome, thus limiting the spread of mitochondrial group II introns into the nascent eukaryotic genome.

## Methods

### Database Construction

Eukaryotic species were manually included in our database by their completeness and position on the most recent tree of life ^49^. Archaea and bacteria were downloaded en masse from Genbank^50^, and ribosomal trees using concatenated universal ribosomal proteins queried from eukaryotic homologues were used to determine their distance from each other. All reference genomes and transcriptomes can be found in the supplemental table, which includes the color codes of all taxa. Bacteria and archaea that were closer than 0.3 substitutions per site overall were too similar, and a species was chosen to represent each cluster that remained this close. Representatives were chosen based on non candidate status and completeness, and in cases where there were more than one that fit said criteria, were selected at random.

### Species Tree Proteins and Concatenations

Universal proteins were queried from the set of universal ribosomal proteins present in eukaryotes as discussed by Eme et al.^49^, and from non ribosomal eukaryotic proteins^51^. All proteins were queried via BLAST v2.15.0+^52^ against a database of curated eukaryotes, archaea and bacteria, with a minimum e-value of 1e-5, and a 60% query match to the original search proteins. Proteins were included if they were able to reach *Eschericia coli*, *Haloferax volcanii*, *Sulfolobus acidocaldarius*, *Thermotoga maritima*, *Bdellovibrio bacteriovorus*, *Hodarchaeoa*, *Nitrososphaera viennensis*, *Thermococcus kodakarensis*, and *Woese archaeon*. These taxa represent a wide distribution across bacteria and archaea, and eukaryotic proteins that were able to reach this deeply in all these taxa were used for concatenation. Only species that included all proteins that passed this first filter were included for concatenation, which included representatives of all archaeal and eukaryotic supergroups. Proteins were aligned with MAFFT v. 7.520^53^ and trimmed with trimal v. 1.4.rev15^54^ using -automated1 parameters. Following this, they were concatenated using awk (scripts are on github), and trees were generated using IQTree v. 2.2.2.7 using automated model finder, which identified 7909 sites, 7730 of which were parsimony informative, and selected LG+I+R10 as the model using AIC and BIC criterion.

### Group II Intron Protein Query and Identification

The query of group II introns was constructed from eukaryotic mitochondrial and chloroplast group II introns from *Arabidopsis thaliana*, and were searched against the same database used above using psiblast^52^ with an e-value of 1e-1. Results were further filtered using HMMer v 3.3.2^55^ on the PFAM protein database (downloaded 9/12/2023). Proteins were included if they had the PFAM motifs Intron_maturas2 or GIIM at a significance of 1e-5. Protein positions in the corresponding nucleotide FASTA files were identified by querying each species’ protein sequence against the nucleotide FASTA using BLAST v.2.15.0+ tblastn. Regions containing the maturase coding sequence were extracted and searched with INFERNAL v. 1.1.5^56^. Valid group II intron RNA structures were identified if they contained Intron_gpII or any group-II-D1D4 motif.

### Alphafold Prediction and Query

Specific group II introns were selected for folding with Alphafold v2^57^ on default conditions. The best predictions were kept for each intron, and compared against Prp8 from *Arabidopsis thaliana* by direct alignment using RCS PDB Pairwise Aligner^58^. Results were directly exported in tabular format.

## Data Availability

The data underlying this study was all obtained from publicly available databases, and can be located in **Supplemental Table 1**. All alignments are freely available on github at https://github.com/jmattic1/GroupIIIntronCodeAvailability/upload/jmattic1-alignments.

## Code Availability

All code used for the informatics in this study are available on github: https://github.com/jmattic1/GroupIIIntronCodeAvailability/tree/jmattic1-scripts

## Inclusion and Ethics

This research was primarily bioinformatic, and did not involve collaborators outside of the University of Maryland.

**Table 1.** Alphafold structural comparisons between Group II introns and *Arabidopsis thaliana* PRP8. Group II introns identified in Figure 2 were assessed for structural similarity to PRP8 (in this case the *Arabidopsis thaliana* PRP8 structure was used). All comparisons were done with the Protein Structure Databank pairwise structure alignment using the top ranked alphafold prediction (**Table 1**). The spliceosomal protein is significantly different from group II intron maturases, with only the 800 central residues being of similar structure. However, within those residues, comparable structures were identified in both the *Bdellovibrio* and *Heimdallarchaeal* Group II intron maturases at similar TM and RMSD scores. Alphafold structures of all proteins were generated using the google colab scripts that are publicly available on default settings. All comparisons were done via the Protein Structural Database Pairwise Structural Analysis, where all columns are compared to the PRP8 gene.

## Acknowledgements

The authors would like to thank Stephen M. Mount, Antony M. Jose, and Sean B. Carroll for helpful comments on the draft manuscript. This research was funded in part by the Gordon and Betty Moore Foundation through Grant GBMF11481 to the University of Maryland.

---

**Supplementary Figure 1.**
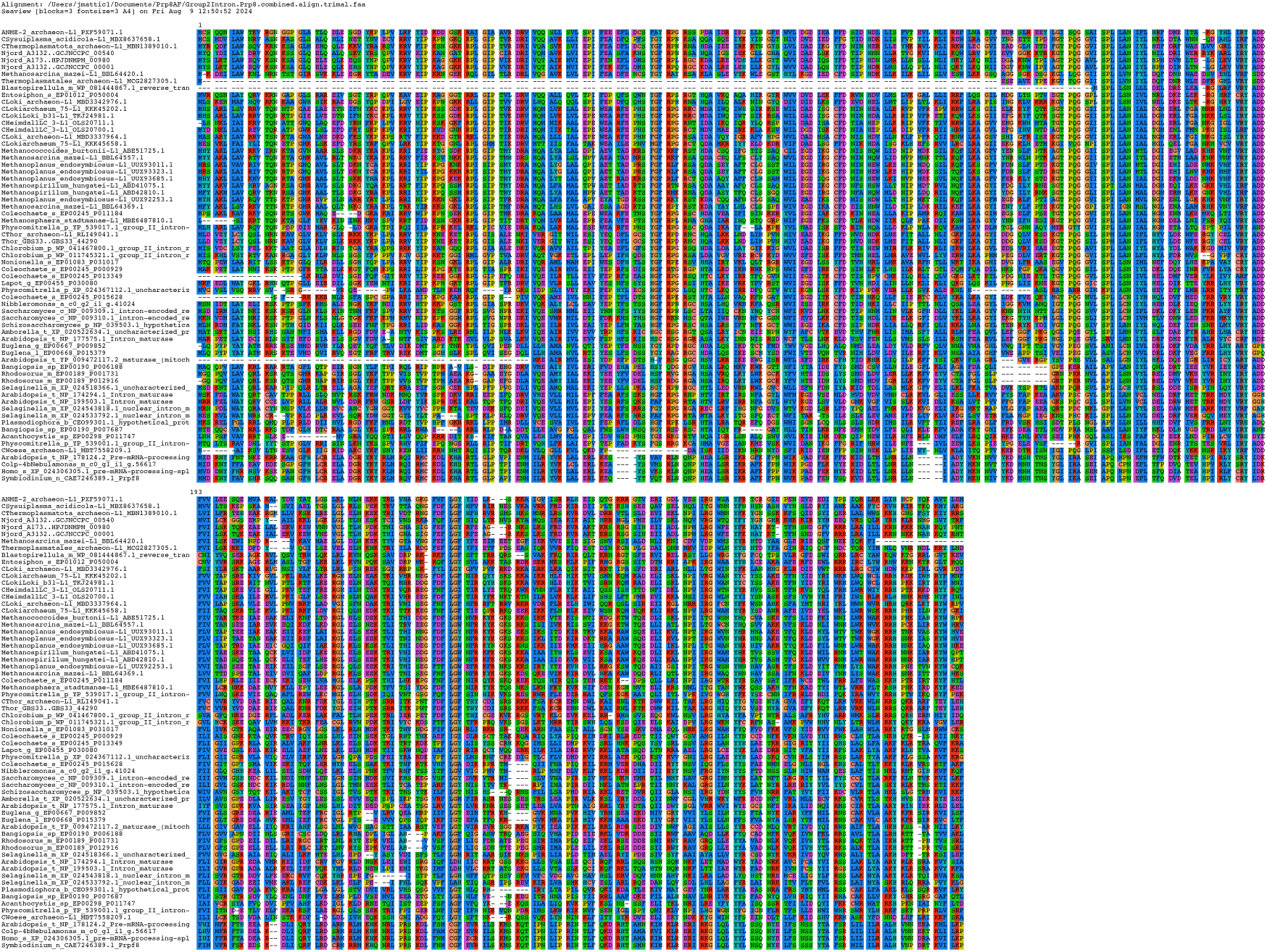
Alignment of eukaryote and archaeal group II introns. Group II intron maturase sequences were aligned with MAFFT and trimmed with trimal with the -gappyout setting. IQTree inferred a maximum likelihood tree from this alignment in Figure 2.

**Supplementary Figure 2.**
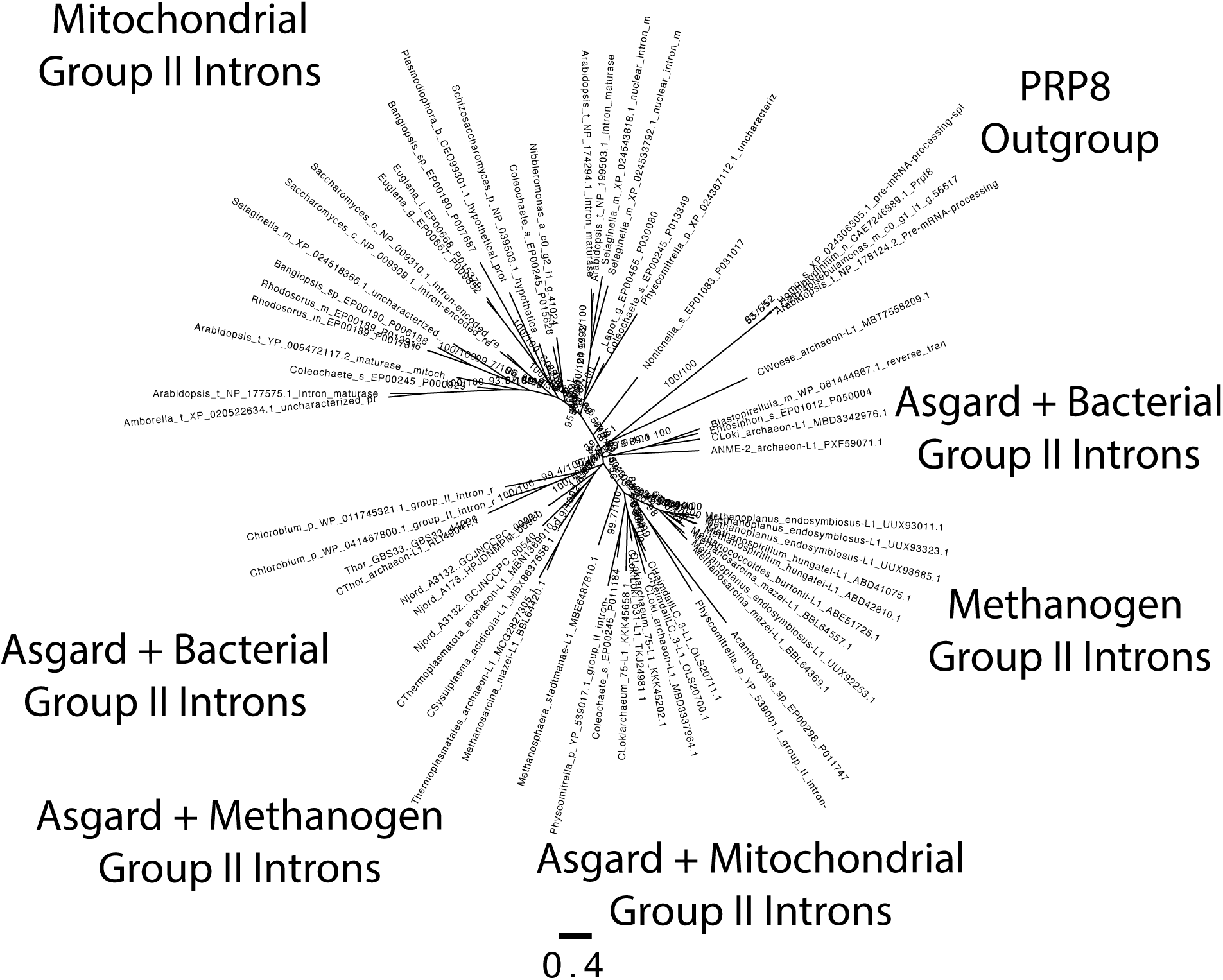
Unrooted tree of eukaryote and archaeal group II introns. Representative group II introns were queried using PSIBLAST against representative eukaryotic, archaeal, bacterial and viral databases. Threshold for inclusion was set at 1e-1 and the search progressed for 10 iterations. Results were then filtered for proteins that contain the characteristic group II intron maturase domain. The resulting tree was left unrooted and indicates that bacterial group II introns appear to be more similar to group II introns already present in the Asgardarchaea than to eukaryotic group II introns. Group II introns were also detected in DPANN, euryarchaeotes and Crenarchaea.

**Supplementary Figure 3.**
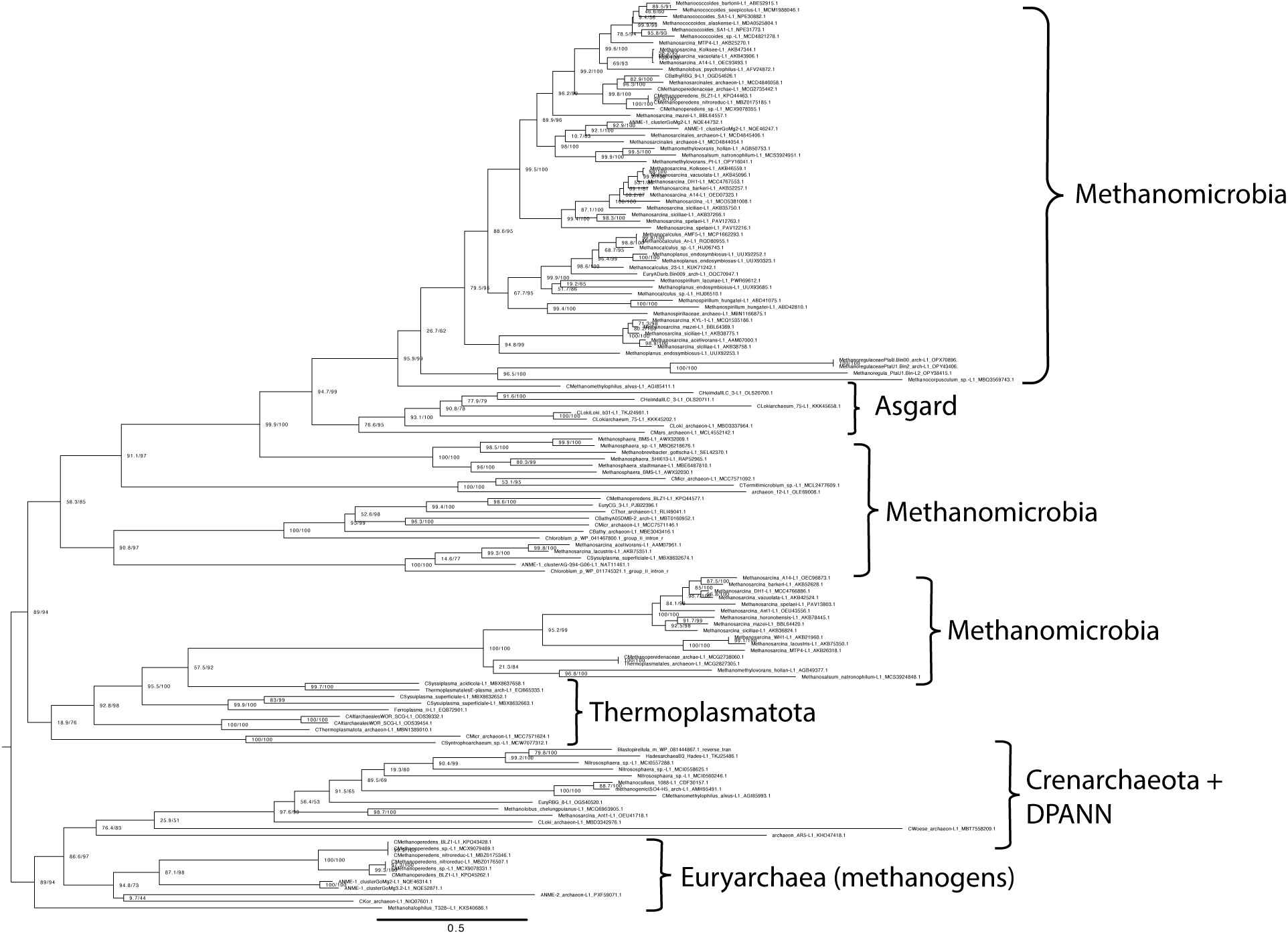
Group II introns across archaea. Representative group II introns were queried using PSI-BLAST against all archaea present in Genbank. Representative group II introns among archaea. Sequences were identified via PsiBLAST at low stringency (1e-1) with ten iterations, then filtered for the maturase domain. The resulting tree was left unrooted and indicates that group II introns are widespread in archaea, including in most major clades of the Asgardarchaea. Group II introns largely form monophyletic clades that correspond to archaeal supergroups, suggesting primarily vertical inheritance of these sequences in the domain.

**Supplementary Figure 3.**
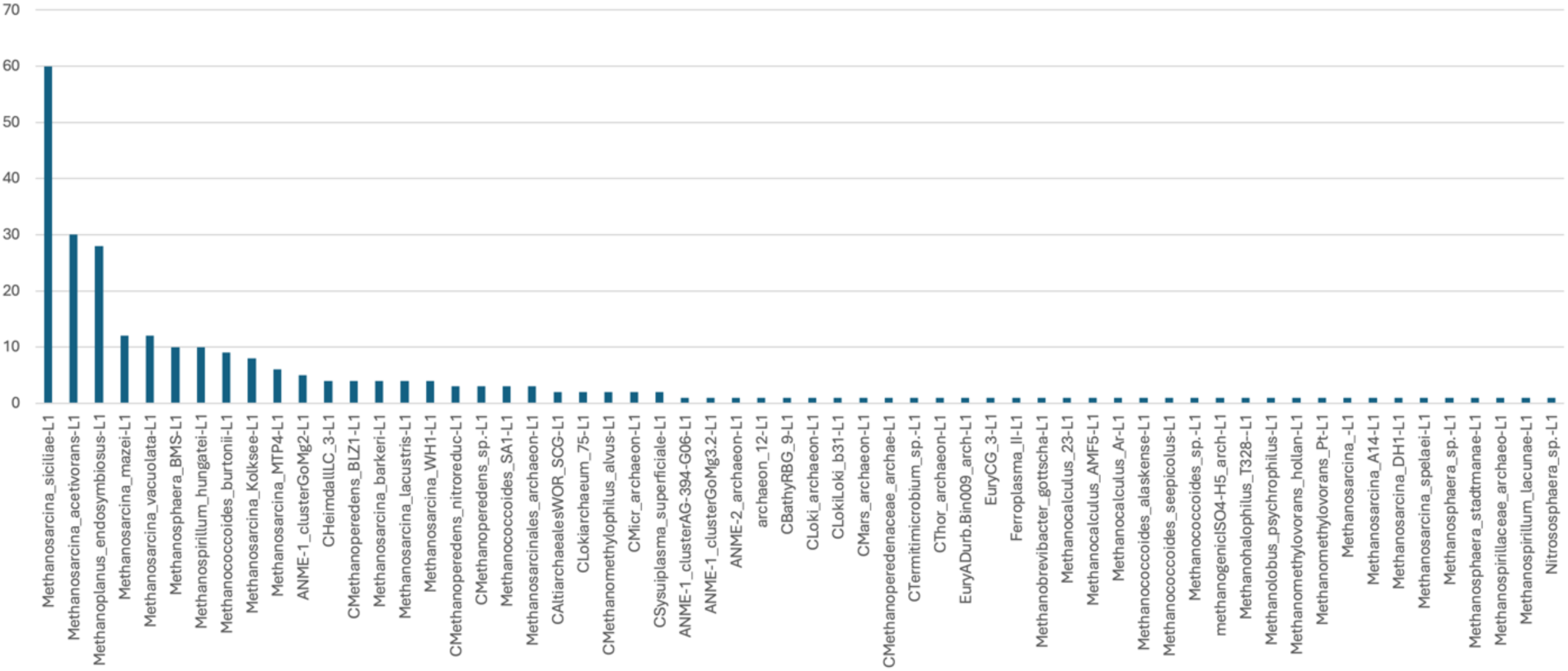
Frequency of group II introns in archaeal genomes. All archaeal genomes that had group II introns with active maturases detected (Figure 3) were downloaded, and the contigs that contained the putative intron were queried with RFAM. All maturases were able to be co-localized to a matching RNA secondary structure hit for the RFAM domain Intron_gpII, indicating the capability for active catalysis. The inferred copy number of group II introns in each genome was calculated based on the number of independent and non-overlapping RNA secondary signatures present in each archaeal genome.

## Footnotes

* Although such a tree is often referred to as a “two domain tree”, a rooted, dichotomously branching tree will always have three basal branches, one solitary and one uniting two other clades. Thus, ignoring gene transfer, the tree of life is expected to be a three domain tree under any circumstances. In this context, the three basal branches appear to be Gracilicutes, Terrabacteria, and Arkarya ^39,40^, or, alternatively, perhaps Bacteria, Euryarchaeota, and “Crenarkarya” or “Protarkarya.” The understanding of the Arkarya tree is currently in rapid development, and this, combined with improved estimates for the timing of diversification of the basal clades should clarify the identity of the three domains.

## Notes

https://github.com/jmattic1/GroupIIIntronCodeAvailability/tree/jmattic1-scripts

### Competing Interest Statement

The authors have declared no competing interest.

